# Calcium alginate entrapped *Eupatorium adenophorum* Sprengel stems powder for Chromium(VI) biosorption in aqueous medium

**DOI:** 10.1101/560680

**Authors:** Mahendra Aryal

## Abstract

A novel biosorbent, *Eupatorium adenophorum* Sprengel was used for biosorption of chromium (VI) from aqueous solutions. Optimum pH, biomass concentration and contact time were estimated at 2.0, 1.0 g/L and 60 min respectively. Data obtained from batch studies confirmed well to the Langmuir model. It was found that the overall biosorption process was best described by pseudo second-order kinetics. The results demonstrated that intraparticle diffusion is not the sole rate-controlling step for the whole sorption process. In binary mixtures, biosorption efficiency of Cr(VI) was not significantly affected by the presence of ions up to 50 mg/L. Thermodynamic study showed that Cr(VI) biosorption on *Eupatorium adenophorum*-alginate beads was spontaneous, exothermic and physisorption in nature. Cr(VI) ions were recovered effectively from *Eupatorium adenophorum-*alginate beads using 0.5 M HNO_3_ solution.

## Introduction

Chromium is the 21^st^ most abundant element in earth crust. It is generally found as trivalent and hexavalent chromium in natural environments. Trivalent chromium occurs naturally in many vegetables, fruits, meat, grains and is often added to vitamins as a dietary supplement, whereas hexavalent chromium, most often generated in the environment by industrial processes and mining of chromium ores [1]. In potable waters, Cr(VI) appears as the most stable species due to the aerobic conditions in the environment. Exposure of Cr(VI) can have damaging effects on the human physiological, neurological and biological systems. Therefore, the maximum allowable limit of Cr(VI) species in drinking water has been established at 0.1 mg/L [2].

Precipitation, coagulation, ion exchange, adsorption, oxidation, reduction, and reverse osmosis techniques have been used to remove chromium from its polluted streams. Many of these approaches are extremely expensive and supposed to be ineffective for practical applications. Due to this reason, there is a need for suitable and cost effective removal technology [3]. The biomaterials such as algae, fungi, yeast, bacteria, agricultural and industrial by-products have been studied as biosorbents for the removal of heavy metals. Biosorption of heavy metals by biomaterials is found to be more effective, particularly for the removal of metal ions at low concentrations [4].

Many researchers have studied various plant biosorbents including *Cicer arientinum* [5], *Polyporus squamosus* [6], *Plataneusorientalis* leaves [7], *Ulmus* leaves [8], *Acacia albida* and *Euclea schimperi* [9], and *helianthus annuus* stem waste [10]. The *Eupatorium adenophorum* Sprengel plant biomass has not yet been used for any biosorption processes. Therefore, biosorbent prepared from this biomass has been used for Cr(VI) removal from aqueous solutions.

The matrices such as calcium alginate, silica, polyacrolylamide, polysulfone, polyethylenimine and polyhydroxoethylmethyacrylate have been used for immobilization of biomaterials [11]. The use of powered biomass for removal of heavy metals on a commercial scale may create problems due to the low density, and poor isolation of solid and liquid phases [3]. Therefore, it needs to consider for immobilization of biomass for heavy metal treatment from industrial effluents to overcome these limitations. The entrapment of biosorbents in the matrix of insoluble Ca-alginate has been used in heavy metals removal. The immobilization of biosorbents in Ca-alginate is soft form of gel particles with strong mechanical strength [11]. The literatures showed that very few efforts have been directed toward the remediation of heavy metals using immobilized biosorbents.

*Eupatorium adenophorum* Sprengel biomass was finally selected to study on various parameters such as contact time, pH, biomass concentration and initial Cr(VI) concentration respectively. Kinetic, isotherm, desorption studies as well as effect of interfering ions on Cr(VI) removal were also investigated. The main aim of this research was the improvement of the sequestration potential of immobilized *Eupatorium adenophorum* Sprengel-alginate beads for removing Cr(VI) ions from aqueous solutions.

## Materials and methods

### Preparation of stock metal solutions

Stock metal solutions were prepared by dissolving appropriate amounts of K_2_Cr_2_O_7_ (Analytical grade) in distilled water to obtain the standard solutions of 1000 mg/L of Cr(VI) ions. The desired metal solutions were prepared by the dilution of the stock standard solution.

### Collection and preparation of biosorbent

*Eupatorium adenophorum* Sprengel species were harvested from Beshishahar Municipality-9, Gandaki Province, Nepal. It is locally known as Banmara, since it is a forest killer and widely spreading as weed in Nepal. *Eupatorium adenophorum* Sprengel stems were first cut into small pieces, dried in sunlight until all the moisture was evaporated and then converted into powdered form. After being powdered, it was treated with 1 M HCl in order to remove the impurities. It was then washed with distilled water as long as the pH of the washing solution becomes in neutral range. After washing, moisture content of biomass was determined by drying a pre-weighted amount in an oven at 100 °C for 24 h.

### Immobilization of biosorbent

Calcium alginate was used as an immobilizing agent for *Eupatorium adenophorum* Sprengel stems powder. Two percentage of sodium alginate was first dissolved in distilled water and it was mixed with calculated amount of dried *Eupatorium adenophorum* Sprengel powder. *Eupatorium adenophorum*-alginate mixture was then dropped by a syringe into the 5% CaCl_2_ solution placed on magnetic stirrer held around 5-8 °C. The beads were hardened in this solution for 24 h in refrigerator at 4 °C [3]. The beads were washed with distilled water for several times to remove excess of calcium ions and untrapped *Eupatorium adenophorum* Sprengel biomass particles. Thus obtained beads were then air dried. It was found that 1.0 g of alginated beads contain 0.052 g of dry biomass. The immobilized biomass was used for further biosorption experiments.

### Biosorption experiments

Biosorption experiments were carried out in 50 ml conical flasks. Initial Cr(VI) concentration of 10 mg/L was used in order to determine the optimum pH, contact time and biomass concentration. The sorption of Cr(VI) ions on *Eupatorium adenophorum* Sprengel*-*alginate beads at varying initial pH values from 1.0 to 7.0, biomass concentration from 1.0 to 7.0, and sorption time from 0 to 130 min were performed. The pH was initially altered by the addition of either 0.1 M NaOH or 0.1 M HNO_3_ to the biosorption systems. Equilibrium isotherm experiments were carried out at various temperatures of 20, 30 and 40°C with initial concentration of Cr(VI) ions varied between 10 and 300 mg/L at pH 2.0, biomass concentration 1.0 g/L and contact time 60 min. respectively. The samples were collected at predetermined time intervals. All experiments were performed in triplicate and the mean values were used in the data analysis. The Cr(VI) concentration was determined at 540 nm followed by complex formation with 1,5-diphenylcarbazide using a spectrophotometer [12].

### Effect of interfering ions on Cr(VI) biosorption

Reference solutions of SO_4_^-2^, Cl^-^, CO_3_^-2^, Mg^+2^, Ca^+2^, Fe^+3^, Zn^+2^, Cd^+2^, Cu^+2^ and Ni^+2^ were prepared from Na_2_SO_4_, NaCl, Na_2_CO_3_, MgSO_4_.7H_2_O, CaCl_2_.2H_2_O, FeCl_3_.6H_2_O, ZnSO_4_.7H2O, Cd(NO_3_)_2_.4H_2_O, CuSO_4_.5H_2_O and NiSO_4_.6H_2_O respectively. The effect of interfering ions ranging from 5 to 50 mg/L on Cr(VI) biosorption at 10 mg/L was carried using the same procedures previously adopted in biosorption experiments. Another series of competitive experiments was also conducted with all mixed ionic systems at concentrations ranging from 5 to 25 mg/ L

### Desorption studies

Biosorption experiments were first conducted with initial Cr(VI) concentrations of 250 mg/L at optimum conditions of pH, biomass concentration and contact time respectively. After sorption of Cr(VI) ions, *Eupatorium adenophorum*-alginate beads were collected carefully and washed with distilled water. The Cr(VI) ions-sorbed beads were dried in oven at 60 °C for 24 h and then again suspended in 30 ml of 0.5 M HNO_3_ solution at room temperature for 2 h. Desorption percentage was calculated from the following equation;

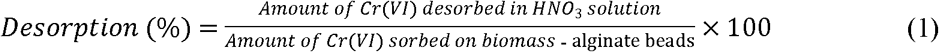

### Data analyses

The amount of Cr(VI) ions sorbed by the *Eupatorium adenophorum*-alginate beads is given by the following equation;

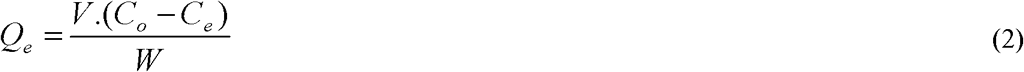

Where *Q*_e_ is the amount of Cr(VI) sorbed by the biomass (mg/g) at equilibrium, *C*_o_ is the initial concentration of Cr(VI) (mg/L), *C*_e_ is the concentration of Cr(VI) at equilibrium (mg/L), *V* is the initial volume of Cr(VI) solution (L), and *W* is the mass of the sorbent (g).

### Langmuir-Hinshelwood (L-H) kinetic model

The Langmuir-Hinshelwood (L-H) kinetic model was used to describe the adsorption kinetics of metal ions on biomass [13]. The linear form of this model can be represented as;

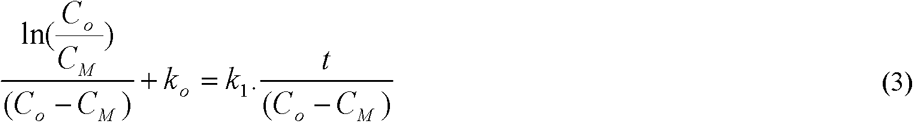

Where *k*_o_ is the reaction rate constant of L-H model (L/mol), *k*_1_ is reaction rate constant (min^-1^), and *C*_*M*_ is metal concentration in solution (mol/L). The plot between [ln(*C*_o_/*C*_M_)]/(*C*_o_-*C*_M_) and *t*/(*Co*-*C*_*M*_) gives a straight line, where the slope is *k*_1_ (min^-1^) and the intercept is *k*_o_ (L/mol) respectively. *k*_*o*_ is similar to a zero order rate constant that represents the initial sorption rate and *k*_*1*_ is first order rate constant after sorption reaches its maximum.

### Avrami kinetic model

The Avrami kinetic model determines the fractionary kinetic orders and some kinetic parameters reflecting possible changes in the sorption rates as function of the initial concentration and sorption time [14]. The linearized form of this equation is;

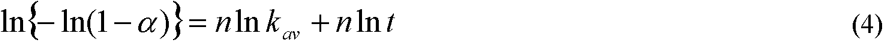

Where *α* is an adsorption fraction (*Q*_t_/*Q*_e_) at time *t, k*_*av*_ is the Avrami kinetic constant (min^-1^) and *n* is a fractionary reaction order. The values of *n* and *k*_*av*_ can be determined from slope and intercept of a plot of ln{−ln(1 − α)} between ln*t* respectively.

### Lagergren pseudo-first order kinetic model

Pseudo first-order kinetic model considers that the rate of occupation of sorption sites is proportional to the number of unoccupied sites [15]. The linear form of pseudo first-order kinetic equation can be written as follows;

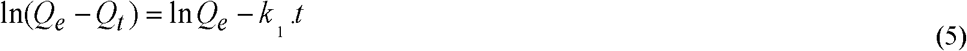

Where *k*_1_ is the first-order rate constant (min^-1^) respectively.

### Ho pseudo-second order kinetic model

Pseudo second-order kinetic model assumes that the rate of occupation of sorption sites is proportional to the square of the number of unoccupied sites [16]. The linear form of this model is given by following equation;

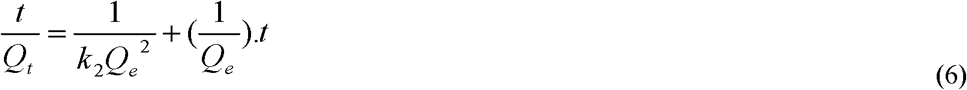

Furthermore, equation (6) can be written as:

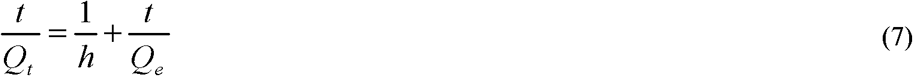

The plot of *t*/*Qt* versus *t* gives a straight line, whose slope and intercept gives *k*_2_ or *h*, and *Q*_*e*_ respectively.

### Ritchie pseudo second-order kinetic model

The basic assumptions of Ritchie second-order kinetic model are; one sorbate is sorbed on two binding sites and rate of sorption depends solely on the fraction of the sites [17]. The linear form of this kinetic model can be written as;

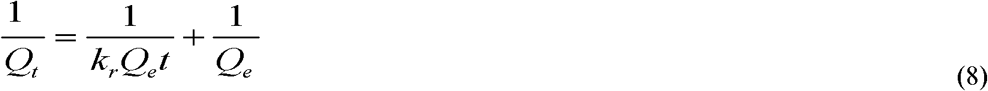

Where *k*_*r*_ is the rate constant of the Ritchie-second-order kinetic model (1/min) and can be obtained from the slope of a line between 1/*Q*_*t*_ vs 1/*t*.

### Sobkowsk and Czerwinski pseudo-second order kinetic model

This model assumes that first order reaction can be applied for lower surface concentrations of solid and the second order for higher surface concentrations [18].

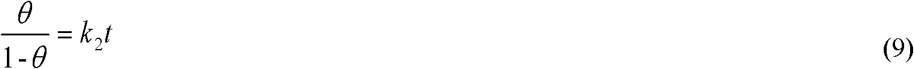

Where *θ* = *Q*_t_/*Q*_*e*_, is the fraction of surface sites that are occupied by an adsorbed solute particles. The second order rate constant *k*_*2*_ (min^-1^) can be evaluated from the plot of *θ* /(1-*θ*) versus time *t*.

### Blanachard pseudo-second order kinetic model

Blanachard et al. [19] proposed a second order rate equation similar to that of Ritchie’s model for the exchange reaction of divalent metallic ions onto NH_4_^+^ ions in fixed zeolite particles. The linearized form of Blanachard’s second order kinetics is;

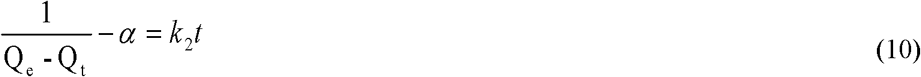

The rate constant, *k*_*2*_ (g/mg.min) can be obtained from the slope of a plot between 1/(*Q*_*e*_-*Q*_t_) versus time *t*.

### Elovich kinetic model

The Elovich chemisorption model is a rate equation based on the adsorption capacity and sorption data [20]. The linear equation is;

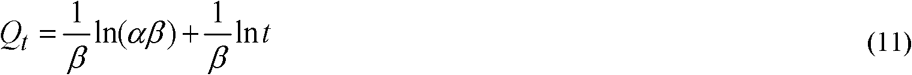

Where *α* is the initial sorption rate constant (mg/g·min), and *β* is related to the extent of surface coverage and activation energy for chemisorptions (g/min). The values of *β* and *α* can be obtained from slope and intercept linear relationship of ln*t* verses *Q*_*t*_.

### Intraparticle diffusion kinetic model

Intraparticle diffusion model explains the diffusion mechanism of the sorption process [21].

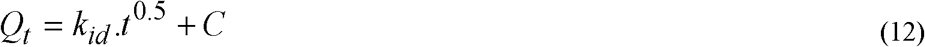

Where *k*_*id*_ is the initial rate of the intraparticle diffusion (mg/g min^0.5^) and C is the intercept of the slope. The values of *k*_*id*_ and *C* can be obtained from the slope and intercept of the linear plots of *Q*_*t*_ versus *t*^*0.5*^.

### Langmuir isotherm model

According to the Langmuir isotherm model, there are a finite number of binding sites, which are homogeneously distributed over the sorbent surface, having the same affinity for sorption of a single molecular layer and there is no interaction between sorbed metal ions [22]. The linear form of Langmuir model is given by following equation;

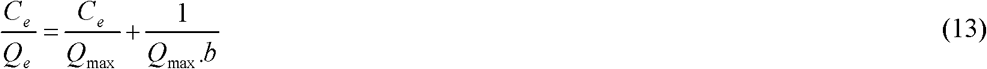

Where *Q*_max_ is the maximum uptake capacity corresponding to site saturation (mg/g) and *b* is the biomass metal binding affinity (L/mg). The adsorption feasibility can be evaluated by the Langmuir isotherm separation factor (*R*_*L*_). It is calculated using equation (14);

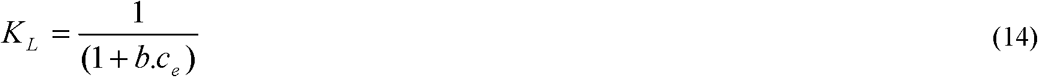

The value of *K*_*L*_ indicates the type of the isotherm to be unfavourable (*K*_*L*_ > 1), linear (*K*_*L*_ = 1), favorable (0 < *K*_*L*_ < 1) or irreversible (*K*_*L*_ = 0).

### Scatchard plot analysis

Scatchard plot explains the number of different types of sites present and their relative affinity for metal ions. The linear plot indicates the single or distinct types of binding sites are present, which is indicative of monolayer coverage [23]. The equilibrium data can be further fitted with Scatchard analysis and its linear form can be written as;

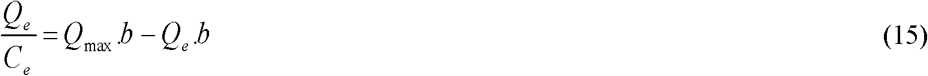

Where *Q*_*max*_ and *b* can be obtained from the slope and intercept of a plot between *Q*_*e*_/*C*_*e*_ and *Q*_*e*_ respectively.

### Freundlich isotherm model

Freundlich isotherm model indicates that the sorption energy of a metal binding to a site on sorbent depends on whether the adjacent sites are already occupied or not. It also explains about the multilayer sorption of metal ions on heterogeneous biomass surface [24]. The linear form of Freundlich equation can be expressed as;

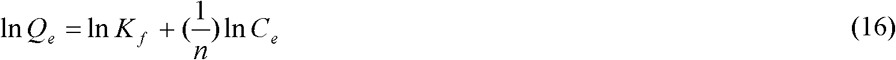

Where, *K*_f_ and *n* are the constants describing sorption capacity and intensity respectively.

### Gin isotherm model

Gin et al., [25] proposed a simple equilibrium isotherm model, which is defined by following linear equation;

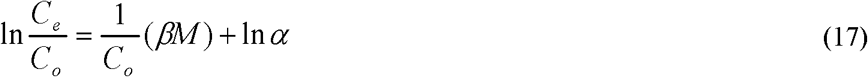

Where *α* and *β* are the parameters of this model. The value of *β* and *α* can be evaluated from the slope and intercept of a plot between ln (*C*_*e*_/*C*_*o*_) and 1/*Co*.

### Temkin isotherm model

This model assumes that the heat of sorption of all molecules in a layer decreases linearly due to the adosorbant-sorbate interaction and that adsorption is characterized by the uniform distribution of binding energies upto some maximum binding energy and can be written in the following linear form [26];

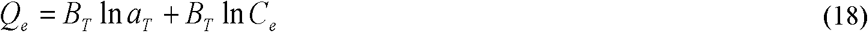

Where;

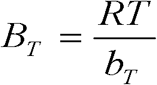

Where *T* is the absolute temperature (K) and *R* is the universal gas constant (kJ/mol). The constant *b*_*T*_ is related to the heat of adsorption (kJ/mol), and *a*_*T*_ (L/mol) is equilibrium binding constant corresponding to the maximum binding energy.

### Dubinin-Radushkeich (D-R) isotherm model

Dubinin-Radushkeich model represents the heterogeneity of the surface energies [27]. This model can be written in the following linear form;

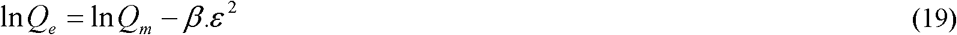

Where;

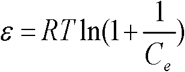

Where *Q*_*e*_ is the amount of metal ions sorbed by biosorbent (mol/g) and *C*_*e*_ in the concentration in equilibrium (mol/L). The constant (*β*), adsorption energy (kJ^2^/mol^2^) and *Q*_*m*_, maximum adsorption capacity (mol/g) can be obtained from slope and intercept of the plot of ln*Q*_*e*_ against *ε*^*2*^. The mean sorption energy (kJ/mol) can be written as:

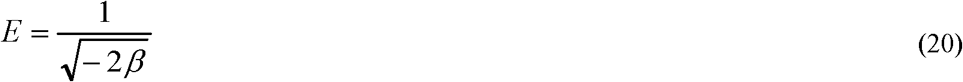

If the magnitude of *E* is in the range of 8 and 16 kJ/mol, sorption process is supposed to be chemisorption, whereas if it is *E* ≤ 8 kJ/mol, physisorption is the predominant mechanism.

### Flory-Huggins isotherm model

The Flory-Huggins isotherm describes the degree of surface coverage characteristics of adsorbate onto adsorbent and also expresses the feasibility and spontaneous nature of an adsorption process [4]. The Flory-Huggins model can be written by following linear equation;

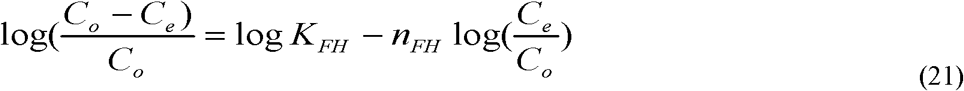

Where, *K*_FH_ is the Flory–Huggins model equilibrium constant (L/mg) and *n*_FH_ the Flory–Huggins model exponent. The Gibbs free energy (Δ*G°*) can be calculated from the following equation;

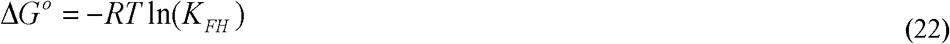

The value of Δ*G°* may reflect, whether the biosorption process proceeds with exothermic or endothermic reaction.

### Hill-der Boer isotherm model

The Hill-der Boer isotherm equation was originally developed to describe the gas-solid phase adsorption to characterize the interactions of adsorbate-adsorbate in the solid phase and adsorbate-adsorbent at the liquid–solid interface [28]. The Hill-der Boer equation can be expressed as;

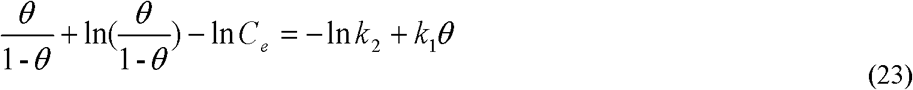

Where *θ* = (1–*C*_*e*_/*C*_*o*_) is the surface coverage fraction, *k*_1_ (dimensionless) and *k*_2_ (mg/L) are constants respectively. The larger the value of *k*_1_, the stronger interactions of adsorbate-adsorbate in the solid phase, while the smaller the value of *k*_2_, the stronger interactions of adsorbate-adsorbent at the liquid-solid interface.

### Halsey isotherm model

Multilayer sorption is generally described by Hasely isotherm model [29]. The linear form of this model is;

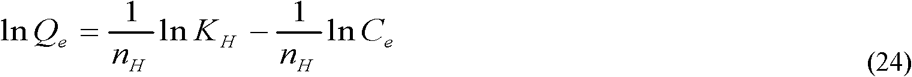

The values of *n*_*H*_ and *K*_*H*_ can be calculated from the slope and intercept of a plot between ln*Q*_*e*_ and ln*C*_*e*_.

### Fundamental thermodynamic parameters

The standard Gibbs free energy change (Δ*G°*) can be also expressed in terms of Langmuir equilibrium constant, *b* (L/mg). Langmuir equilibrium constant, *b* (L/mg) is first calculated from Langmuir equation at different temperatures. In order to find the standard Gibbs free energy change (Δ*G°*), the calculated values of ‘*b*’ (L/mg) should be converted into L/mol [30]. The Gibbs free energy change (Δ*G°*) can be determined using the following equation;

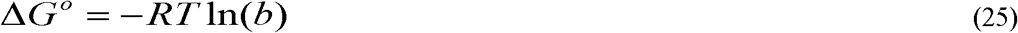

The van’t Hoff equation can be used to determine the enthalpy (Δ*H°*) and entropy change of biosorption (Δ*S°*) as a function of temperature.

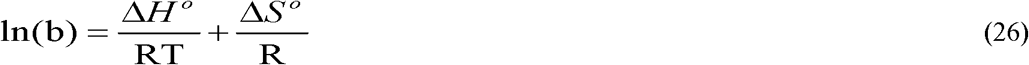

Δ*H°* and Δ*S°* can be obtained from the slope and intercept of a plot of *lnb* versus 1/*T*.

### Activation energy

The activation energy (*E*_a_) is an energy level that must be achieved over and above the initial energy level of a substance in order for a reaction to occur. The greater the activation energy gives the slower reaction rate. The pseudo-second order rate constant can be expressed in Arrhenius equation [31];

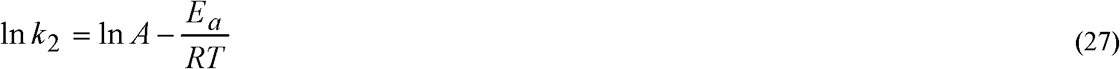

Where *A* is the frequency factor (mg/g min). A plot of ln*k*_*2*_ versus 1/*T* gives a straight line and the value of activation energy (*E*_*a*_) can be determined from its slope.

### Isosteric heat

The heat of adsorption determined for a constant amount of metal ions adsorbed is known as the isosteric heat of adsorption (*ΔH*_*r*_) and its magnitude can be calculated by Clausius–Clapeyron equation;

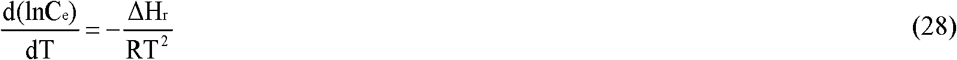

Where *C*_*e*_ is the equilibrium concentration at constant amount of adsorbed Cr(VI) (mol/L) that can be obtained from the biosorption data at different temperatures. Δ*H*_*r*_ can be calculated from the slope of a plot of ln*C*_*e*_ versus 1/*T* [32].

## Results and discussion

### Evaluation of precision and accuracy of Cr(VI) concentrations

The calibration curve was made in the linear range of Cr(VI) concentrations from 0.3 to 2 mg/L with absorbance between 0.159 and 1.137 using a spectrophotometer (Shimadzu UV-160A, Kyoto, Japan). The linear regression result of absorbance against Cr(VI) concentration was A=0.5704C-0.0012 with correlation coefficient of 0.9988. The quality control samples against the number of samples taken are shown by Fig. 1.

**Fig 1.**
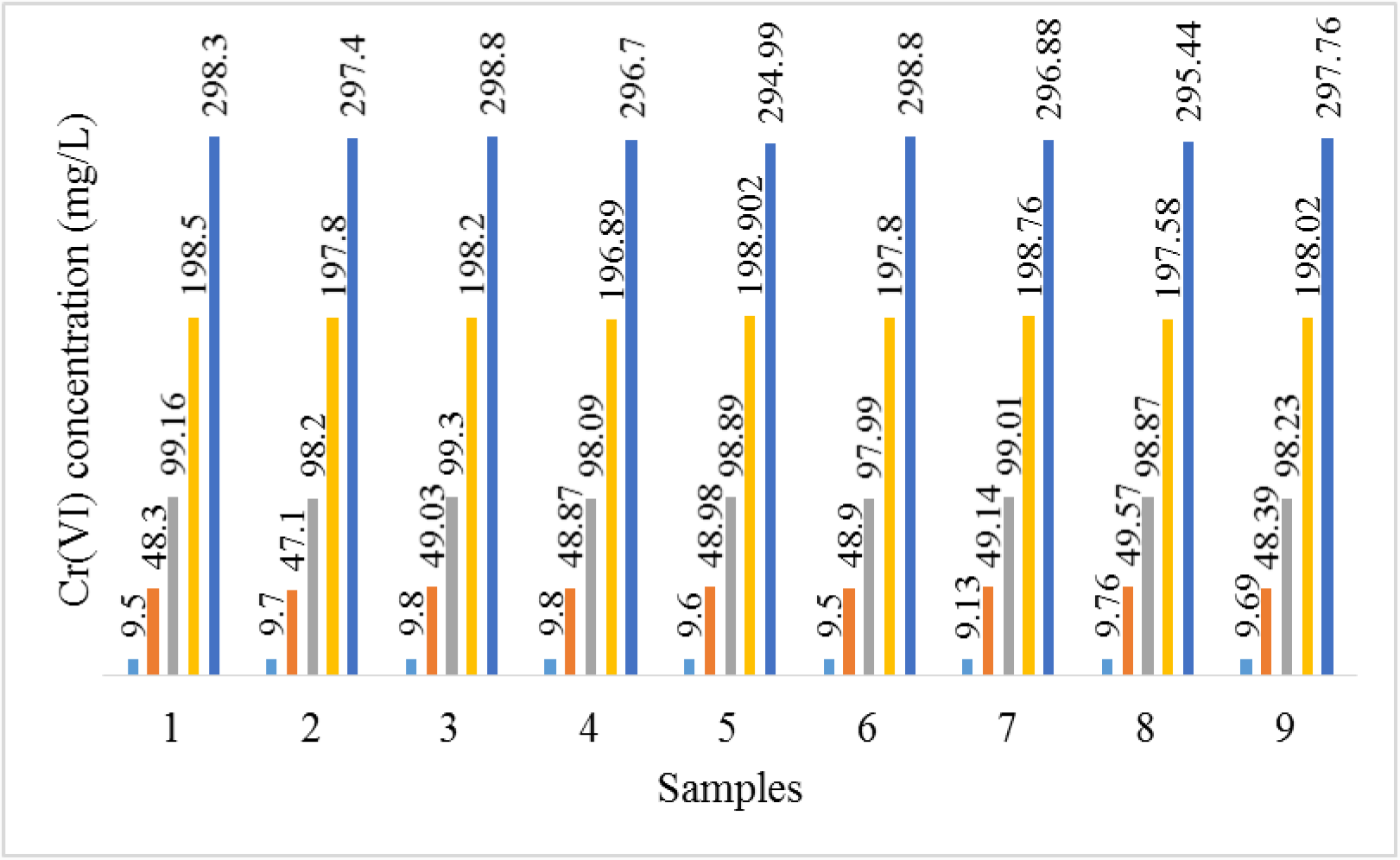
Quality control and number of samples.

**Fig 2.**
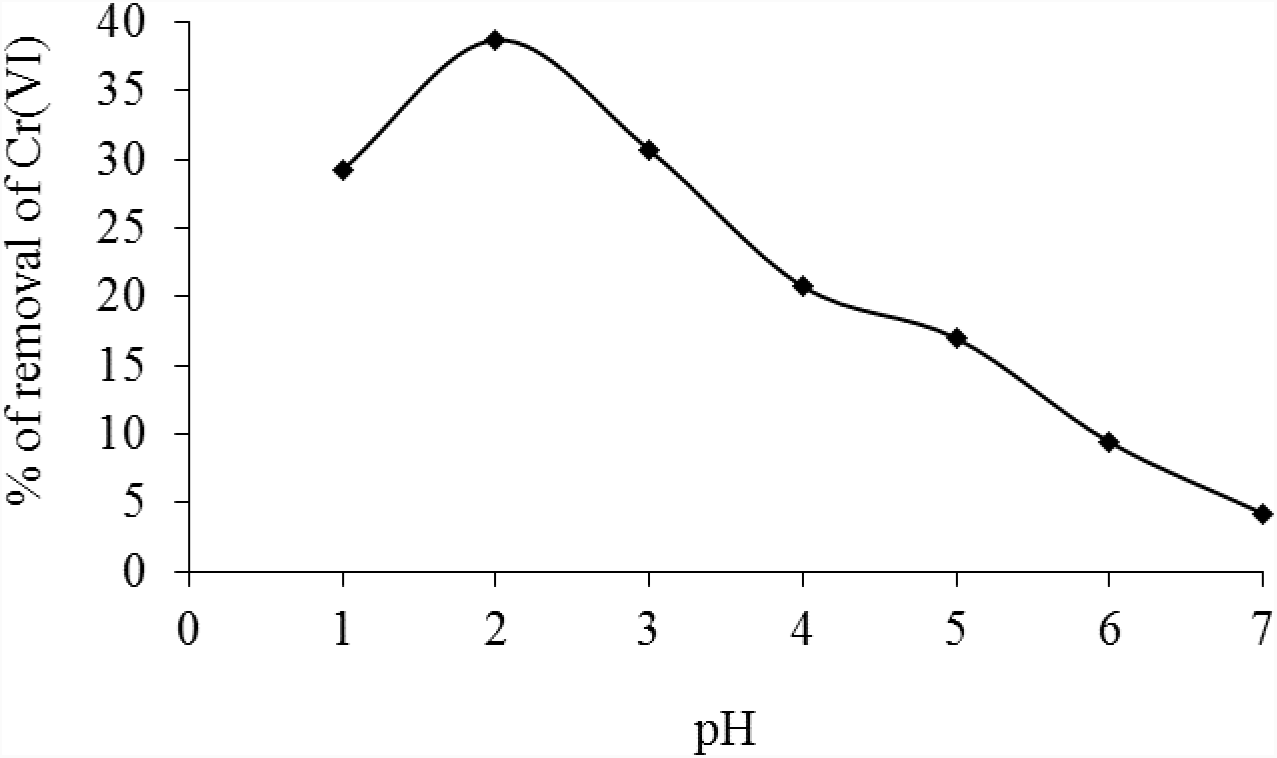
Effect of pH on biosorption of Cr(VI) by *Eupatorium adenophorum–*alginate beads at initial Cr(VI) concentration 10mg/L and biomass concentration 1.0 g/L respectively.

The precision is expressed by relative standard deviation and variance methods. Relative standard deviation is a statistical measurement that describes the spread of data with respect to the mean and the result is expressed as a percentage. RSD and variance are routinely used to assess the variation of sets of data in analytical chemistry. The accuracy is determined in terms of recovery percentage. The precision and accuracy results are presented in Table 1.

**Table 1.** Precision and accuracy data of Cr(VI) analyses

| QC samples (mg/L) | Average (mg/L) | SD | RSD (%) | Variance | Recovery (%) |
| --- | --- | --- | --- | --- | --- |
| 10 | 9.60 | 0.219 | 2.219 | 0.045 | 96.088 |
| 50 | 48.697 | 0.709 | 0.456 | 0.502 | 97.395 |
| 100 | 98.637 | 0.579 | 0.512 | 0.255 | 98.637 |
| 200 | 198.05 | 0.627 | 0.350 | 0.393 | 99.025 |
| 300 | 297.238 | 1.369 | 0.460 | 1.876 | 99.076 |

**Note:** QC; Quality control; SD: Standard deviation; RSD: Relative standard deviation

### Effect of pH

Solution pH is one of the important parameters that significantly influences the sorption of Cr(VI) on sorbent. The effect of pH on Cr(VI) biosorption onto *Eupatorium adenophorum*-alginate beads is shown in Fig.2. The highest Cr(VI) biosorption was determined at pH 2.0. Under acidic conditions, the surface of the biomass becomes highly protonated and favors the sorption of Cr(VI) in the form of HCrO_4_^-^ ions. The decrease in percentage removal of Cr(VI) ions was observed above pH 2.0. This may be due to the gradual increase in negatively charged biomass surface groups and shifting of monovalent HCrO_4_^-^ to divalent Cr_2_O_7_^-2^ and CrO_4_^-2^ ions in aqueous solutions [33].

### Effect of biomass concentration

The effect of biomass concentrations on Cr(VI) sorption is shown in Fig. 3. The results indicated that the percentage removal of Cr(VI) ions was increased with increasing biomass concentrations from 1.0 to 7.0 g/L respectively. The trend of increase in removal capacity is due to the fact that the availability of more binding sites of *Eupatorium adenophorum*-alginate beads for Cr(VI) ions. No significant increment in percentage removal of Cr(VI) ions was observed above biomass concentration of 1.0 g/L (Fig. 3a). This may be due to the strong limitations of Cr(VI) ions mobility in the sorption medium in a fixed volume, leaving some binding sites unsaturated. On the other hand, the uptake capacities rapidly decreased with increase of biomass concentrations from 1.0 to 7.0 g/L respectively (Fig. 3b). The low sorption capacity can be ascribed to the fact that *Eupatorium adenophorum*-alginate beads have a limited number of active sites that would have achieved saturation above a certain biomass concentration [34].

**Fig 3.**
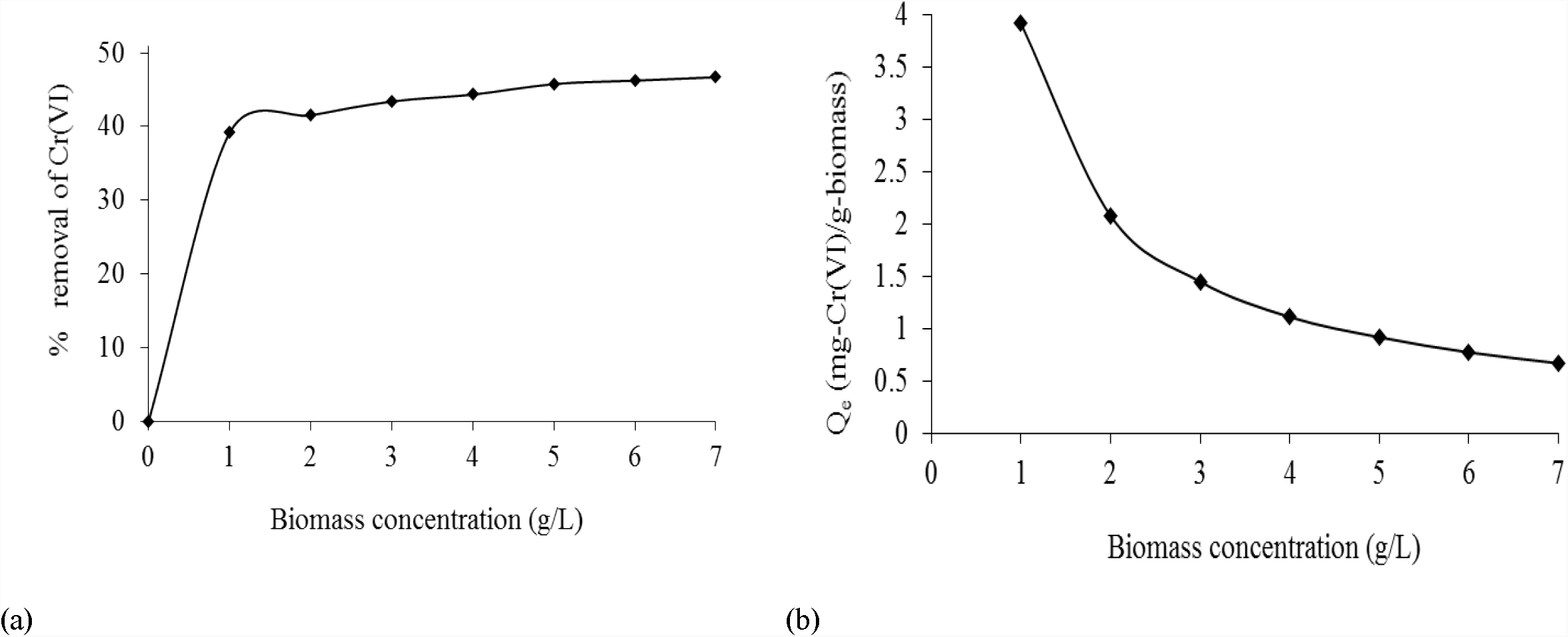
Effect of biomass concentration on biosorption of Cr(VI) by *Eupatorium adenophorum* – alginate beads at initial Cr(VI) concentration 10 mg/L and pH 2.0 respectively.

### Effect of contact time and temperature

The effect of contact time for Cr(VI) biosorption on *Eupatorium adenophorum*-alginate beads at different temperature is shown in Fig. 4. The results indicated that biosorption efficiency increased with increase in contact time rapidly and thereafter proceeded at a lower rate and attained equilibrium condition. The sorption equilibrium time at 60 min might be a result of sorption-desorption processes occurring after saturation of Cr(VI) ions on biomass surface. After 60 min of equilibrium time, the removal efficiency of Cr(VI) was almost constant, suggesting that an equilibrium balance for sorption process [35]. The results also demonstrated that sorption efficiency for Cr(VI) increases with increase in temperature from 20 to 30 °C and this may be due to the increase in collision frequency between Cr(VI) ions and the biomass species. The removal efficiency of Cr(VI) was slightly decreased upon further increase of temperature from 30 to 40 °C, which may be responsible for destruction of some binding sites.

**Fig 4.**
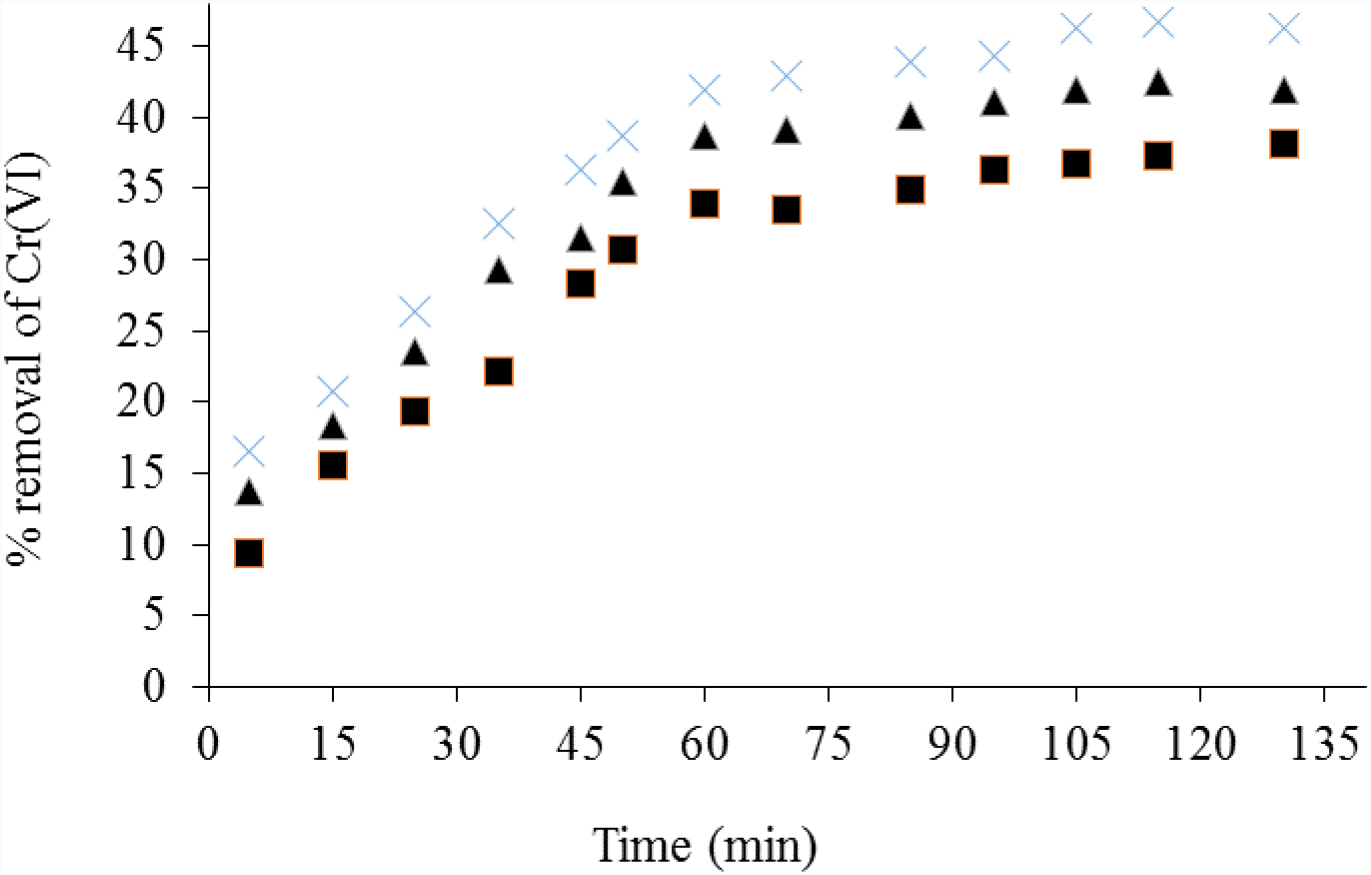
Effect of contact time on on Cr(VI) biosorption by *Eupatorium adenophorum*–alginate beads at initial Cr(VI) concentration 10 mg/L, pH 2.0and biomass concentration 1.0 g/L at 20 (-▄--), 30 (-×-) and 40°C (-▴-) respectively.

### Effect of initial Cr(VI) concentrations

The initial concentration of metal ions in the solution is an important parameter as the metal concentration changes over a broad range in industrial effluents. The effect of initial Cr(VI) concentrations on equilibrium uptake capacity at different temperatures is shown in Fig. 5. It was found that the amount of equilibrium sorption capacity increases with increasing initial Cr(VI) concentrations. At lower initial metal concentrations, sufficient sorption sites are available on biomass surface for biosorption of Cr(VI) ions. At higher concentrations, relatively less available sites induced the reduction in sorption of Cr(VI) ions on *Eupatorium adenophorum*-alginate beads [35].

**Fig 5.**
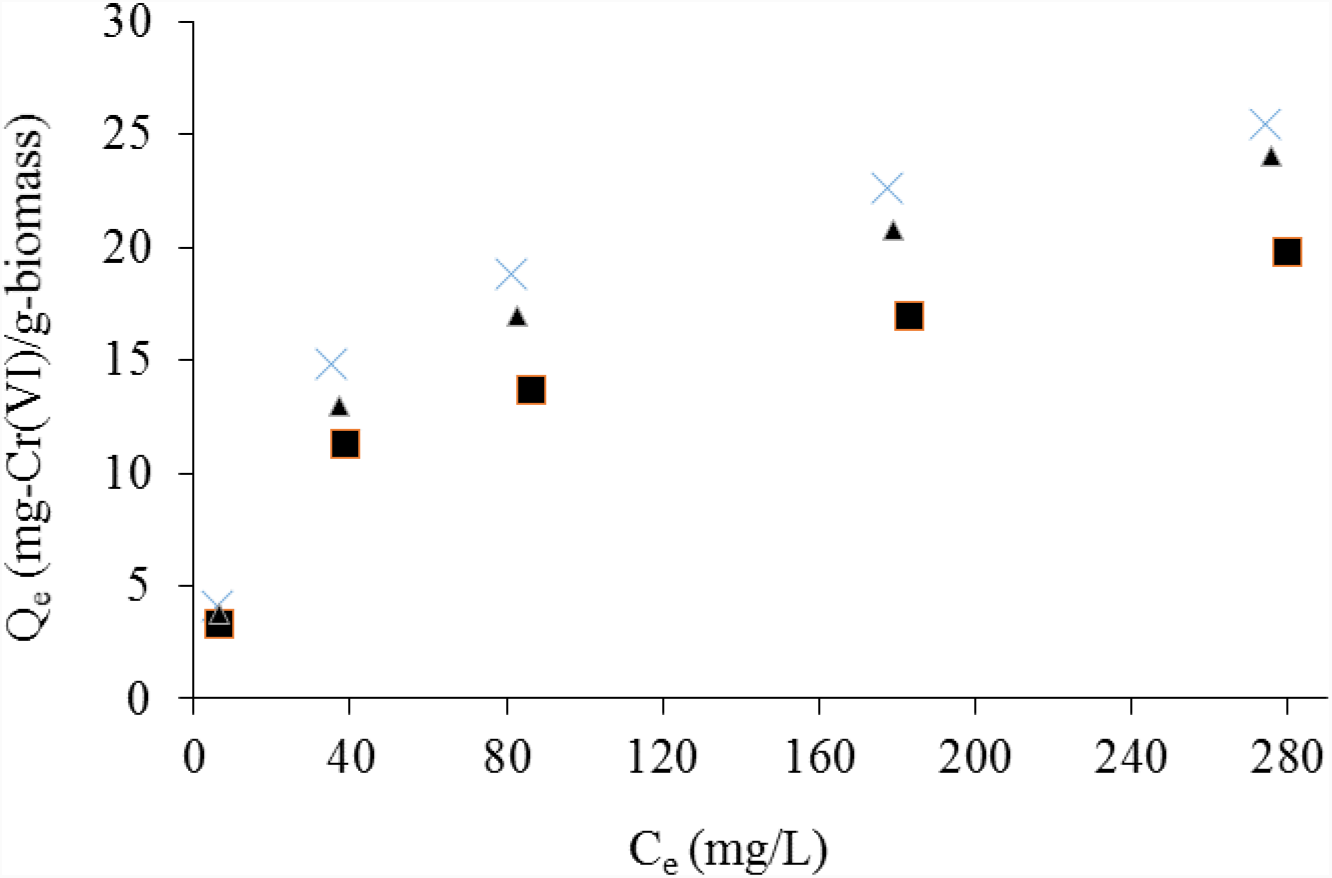
Effect of initial concentration of Cr(VI) ions onto *Eupatorium adenophorum*–alginate beads at initial Cr(VI) concentration from 10 to 300 mg/L, pH 2.0, contact time 60 min and biomass concentration 1.0 g/L at 20 (-▄--), 30 (-×-) and 40°C (-▴-) respectively.

### Determination of kinetic parameters

Kinetic study is important in determining the efficiency of sorption systems. Sorption kinetics expressed as the solute removal rate that controls the residence time of the sorbate in solid-solution interface. Kinetic constants and correlation coefficients of kinetic models are given in Table 2.

**Table 2.** Kinetic parameters of Cr(VI) sorption on *Eupatorium adenophorum*–alginate beads at initial Cr(VI) concentration 10 mg/L, pH 2.0 and biomass concentration 1.0 g/L respectively

| Kinetic model | Parameter | Temperature |  |  |
| --- | --- | --- | --- | --- |
|  |  | 20 °C | 30 °C | 40 °C |
| Langmuir-Hinshelwood | Experimental $Q_e$ (mg/g) | 3.820 | 4.669 | 4.198 |
| | $k_l$ (min<sup>-1</sup>) | 0.008 | 0.001 | 0.009 |
| | $k_0$ (L/mg) | 0.103 | 0.108 | 0.107 |
| | $k_1/k_0$ (mg/L.min) | 0.007 | 0.010 | 0.0083 |
| | $R^2$ | 0.848 | 0.857 | 0.835 |
| Avrami | $n$ | 0.861 | 0.743 | 0.811 |
| | $k_{av}$ (min<sup>-1</sup>) | 0.082 | 0.180 | 0.126 |
| | $R^2$ | 0.961 | 0.936 | 0.933 |
| Lagergren pseudo-first order | $k_1$ (min<sup>-1</sup>) | 0.057 | 0.052 | 0.0529 |
| | $Q_e$ (mg/g) | 6.382 | 5.353 | 5.228 |
| | $R^2$ | 0.877 | 0.931 | 0.921 |
| Ritchie pseudo second-order | $Q_e$(mg/g) | 4.486 | 3.861 | 4.163 |
| | $k_2$ (min<sup>-1</sup>) | 0.105 | 0.060 | 0.088 |
| | $R^2$ | 0.859 | 0.950 | 0.892 |
| Sobkowsk and Czerwinski pseudo second-order | $k_2$ (min<sup>-1</sup>) | 0.302 | 0.260 | 0.397 |
| | $R^2$ | 0.790 | 0.900 | 0.753 |
| Blanachard pseudo second-order | $k_2$ (g/mg min) | 0.079 | 0.0563 | 0.0946 |
| | $\alpha$ (g/mg) | 1.758 | 0.809 | 1.884 |
| | $R^2$ | 0.830 | 0.890 | 0.901 |
| Ho pseudo second-order | $k_2$ (gm/g min) | 0.0079 | 0.0085 | 0.0084 |
| | $h$ (mg/g)/min | 0.161 | 0.259 | 0.220 |
| | $Q_e$ (mg/g) | 4.710 | 5.494 | 5.107 |
| | $R^2$ | 0.980 | 0.988 | 0.987 |
| Elovich | $\alpha$ (mg/g.min) | 2.700 | 5.143 | 3.214 |
| | $\beta$ (g/mg) | 1.054 | 2.313 | 1.537 |
| | $R^2$ | 0.952 | 0.953 | 0.953 |
| Intraparticle diffusion | $k_{id}$ (mg/g min<sup>0.5</sup>) | 0.327 | 0.354 | 0.336 |
| | $C$ (mg/g) | 0.422 | 1.041 | 0.819 |
| | $R^2$ | 0.938 | 0.932 | 0.928 |

The high correlation coefficient value of experimental kinetic data for Langmuir-Hinshelwood (L-H) kinetic model indicates that chemisorption as a predominant mechanism [28]. In our case, low correlation coefficient values indicate that this model could not be used to describe the kinetics of Cr(VI) sorption on *Eupatorium adenophorum-*alginate beads. It further showed that biosorption of Cr(VI) may be physisorption as predominant mechanism [4].

Avrami model suggests possible changes of the sorption mechanism during the sorption process with multiple kinetic orders that alter during the contact of sorbate with sorbent [14]. The observed *R*^2^ values indicated the biosorption of Cr(VI) on *Eupatorium adenophorum-*alginate beads did not follow this model.

The correlation coefficients for the pseudo first-order equation obtained were low and *Q*_e,cal_ values are not in agreement with *Q*_e,exp_ values, suggesting that Cr(VI) biosorption on *Eupatorium adenophorum* biomass is not a pseudo-first order reaction [4].

The regression (*R*^*2*^) values obtained at different temperatures are very high and the adequate fitting of the plots confirmed that the sorption of Cr(VI) by *Eupatorium adenophorum*-alginate beads followed pseudo second order kinetics. It was also found that the experimental, *Q*_e_ values are agreement with the ones obtained from this kinetic model. The rate constant (*k*_2_) and initial sorption rate (*h*) values were higher at 30 °C and these results further revealed that the biosorption process for Cr(VI) species becomes faster at 30 °C [16].

The low correlation coefficient values of Ritchie, Sobkowsk and Czerwinski, and Blanachard pseudo second-order kinetic models showed that these models are unable to explain the experimental kinetic data for biosorption of Cr(VI) on *Eupatorium adenophorum*-alginate beads [4]

The high correlation coefficient values indicated that good agreement of Elovich kinetic model with the experimental kinetic data, suggesting rate-limiting step may be chemisorption process. However, relatively low *R*^*2*^ values at all temperature suggested that biosorption of Cr(VI) species onto *Eupatorium adenophorum*-alginate beads may not chemisorption mechanism [20].

The biosorption process is also studied using the Weber and Morris intraparticle diffusion model. The low correlation coefficient values obtained from intraparticle diffusion model indicated the sorption is not occurring in the pores of *Eupatorium adenophorum*-alginate beads. It was also found that values of intraparticle diffusion rate constant, *k*_*id*_ increase with increasing temperature from 20 to 30 °C and then decreases to 40 °C. This increase in intraparticle diffusion rate constant with increasing temperature may suggest the faster diffusion as well as biosorption process. On the contrary, values of intercept were found to increase from 20 to 30 °C and then decrease to 40 °C. Increase in values of intercept may indicate that increase in thickness of the boundary layer and decrease in chance of the external mass transfer, and hence increase in chance of internal mass transfer. The plots of *Q*_*t*_ versus *t*^*0.5*^ at different temperatures are given in Fig. 6, indicated the invalidation of this model. As it can be seen from these figures, intraparticle diffusion is not the sole rate-controlling step for the whole sorption process, since the plots do not pass through the origin. However, multilinearity indicates that Cr(VI) biosorption may take place in multiple steps. The first linear portions may indicate the boundary layer diffusion of Cr(VI) ions, whereas the second portions suggest the final equilibrium for which the intraparticle diffusion starts to slow down due to extremely low Cr(VI) concentration left in the sorption medium [21]. Therefore, chromium biosorption on *Eupatorium adenophorum*-alginate beads seems to be governed by a two steps mechanism.

### Biosorption isotherms

Adsorption isotherm explains the interaction between sorbate and sorbent and is critical for design of sorption process. The parameters and correlation coefficients of various isotherm models are listed in Table 3.

**Table 3.**
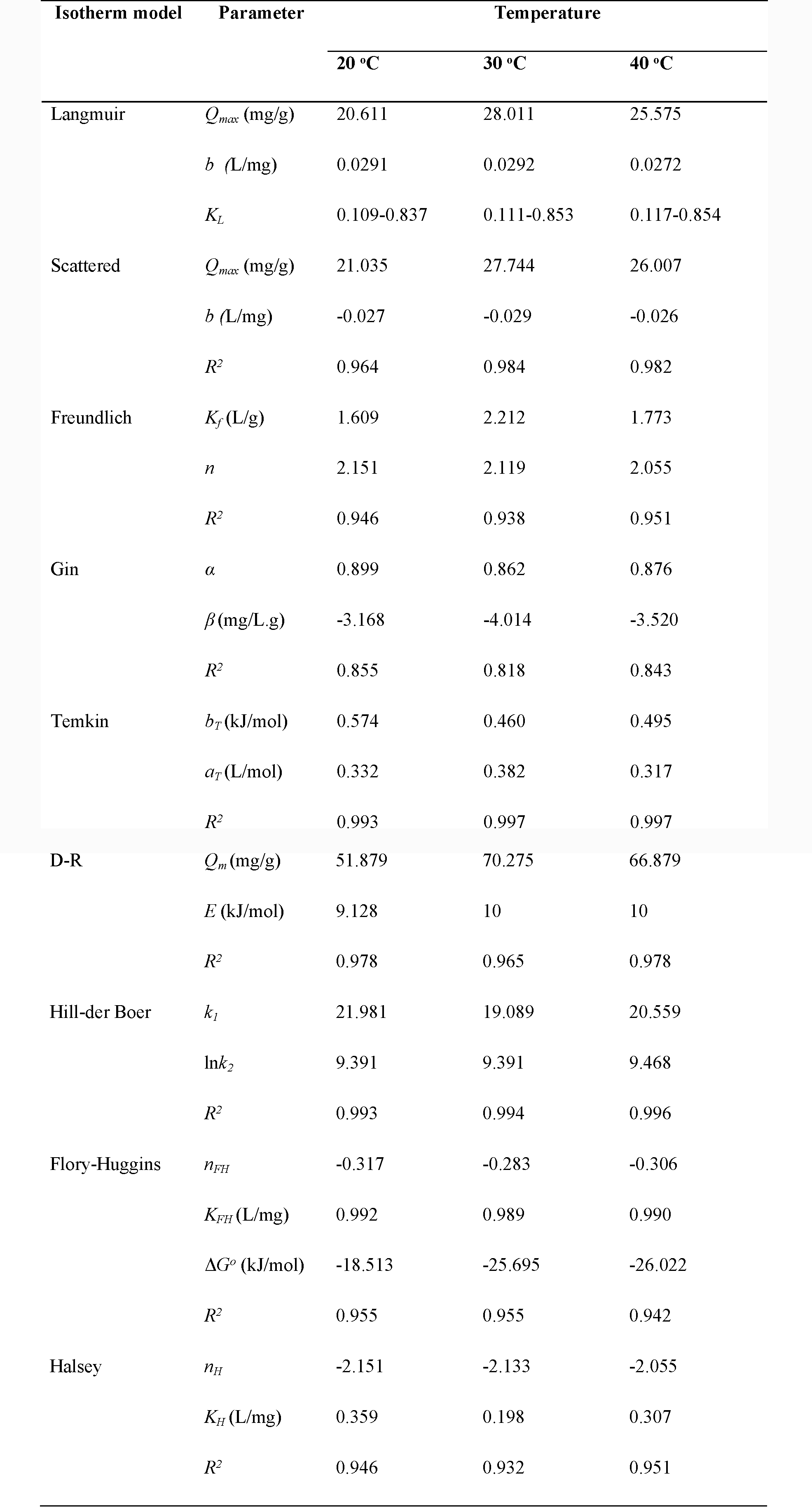
Isotherm parameters of Cr(VI) sorption on *Eupatorium adenophorum*–alginate beads at initial Cr(VI) concentration from 10 to 300 mg/L, pH 2.0 and biomass concentration 1.0 g/L respectively

The application of Langmuir isotherm plots for Cr(VI) biosorption at different temperatures are shown in Fig. 7. The maximum uptake capacity of alginate entrapped *Eupatorium adenophorum* Sprengel biomass for Cr(VI) was found to be at 28.011 mg/g at 30 °C. The high correlation coefficients may be attributed to the homogenous distribution of binding sites on biomass surface and monolayer sorption of Cr(VI) species on *Eupatorium adenophorum* Sprengel biomass surface. In addition, the *R*_*L*_ values found in the range of 0 to 1, indicated that Cr(VI)) biosorption process is favourable for bioorption of Cr(VI). The present alginated *Eupatorium adenophorum* Sprengel biomass has been compared with different plant biosorbents reported in the literature as shown in Table 4. Direct comparison of the performance of biosorbent with other biosorbents might be difficult, since different experimental conditions have been applied.

**Table 4.** Comparison of Cr(VI) uptake capacity with other plant biosorbents

| Biosorbent | $Q_{max}$ (mg/g) | Reference |
| --- | --- | --- |
| *Cicer arietinum* | 72.16 | [5] |
| *Polyporus squamosus* | 31.2 | [6] |
| *Eupatorium adenophorum Sprengel* | 18.587 | Present study |
| Neem leaves | 10.40 | [36] |
| *Eichhorniacrassipes* | 6.0 | [37] |
| *Plataneusorientalis* leaves | 5.01 | [7] |
| *Euclea schimperi* | 3.946 | [9] |
| *Acacia albida* | 2.983 | [9] |
| *Ulmus* leaves | 0.9 | [8] |

**Fig 7.**
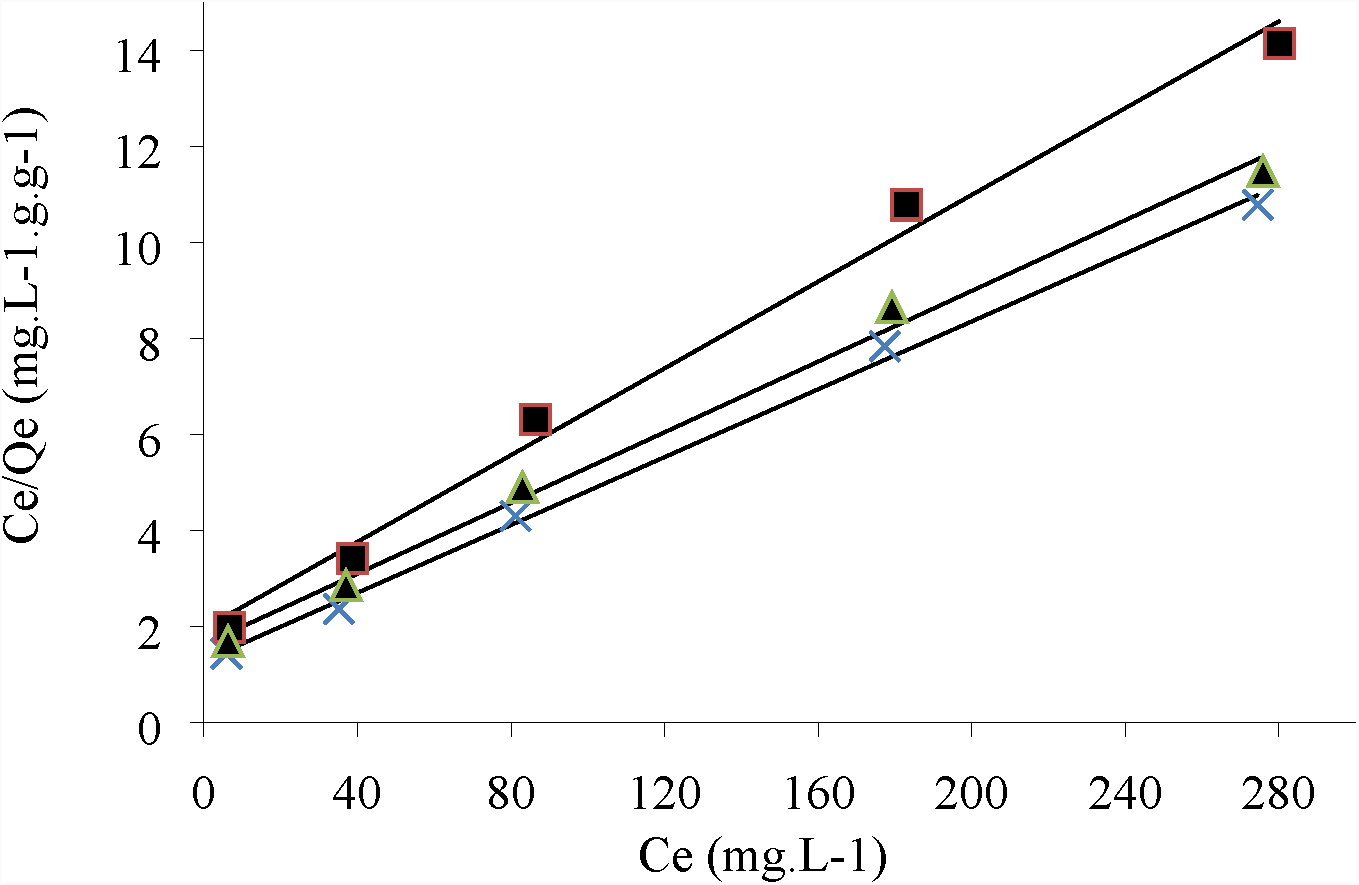
Langmuir isotherm for Cr(VI) ions onto *Eupatorium adenophorum*–alginate beads at initial Cr(VI) concentration from 10 to 300 mg/L, pH 2.0, contact time 60 min and biomass concentration 1.0 g/L at 20 (-▄--), 30 (-×-) and 40°C (-▴-) respectively.

The Scatchard plots for Cr(VI) biosorption on *Eupatorium adenophorum* Sprengel biomass at different temperatures are given in Fig. 8. The linear plots at different temperatures confirmed that the binding sites of this biosorbent exhibits the same affinity towards Cr(VI) ions [23].

**Fig 8.**
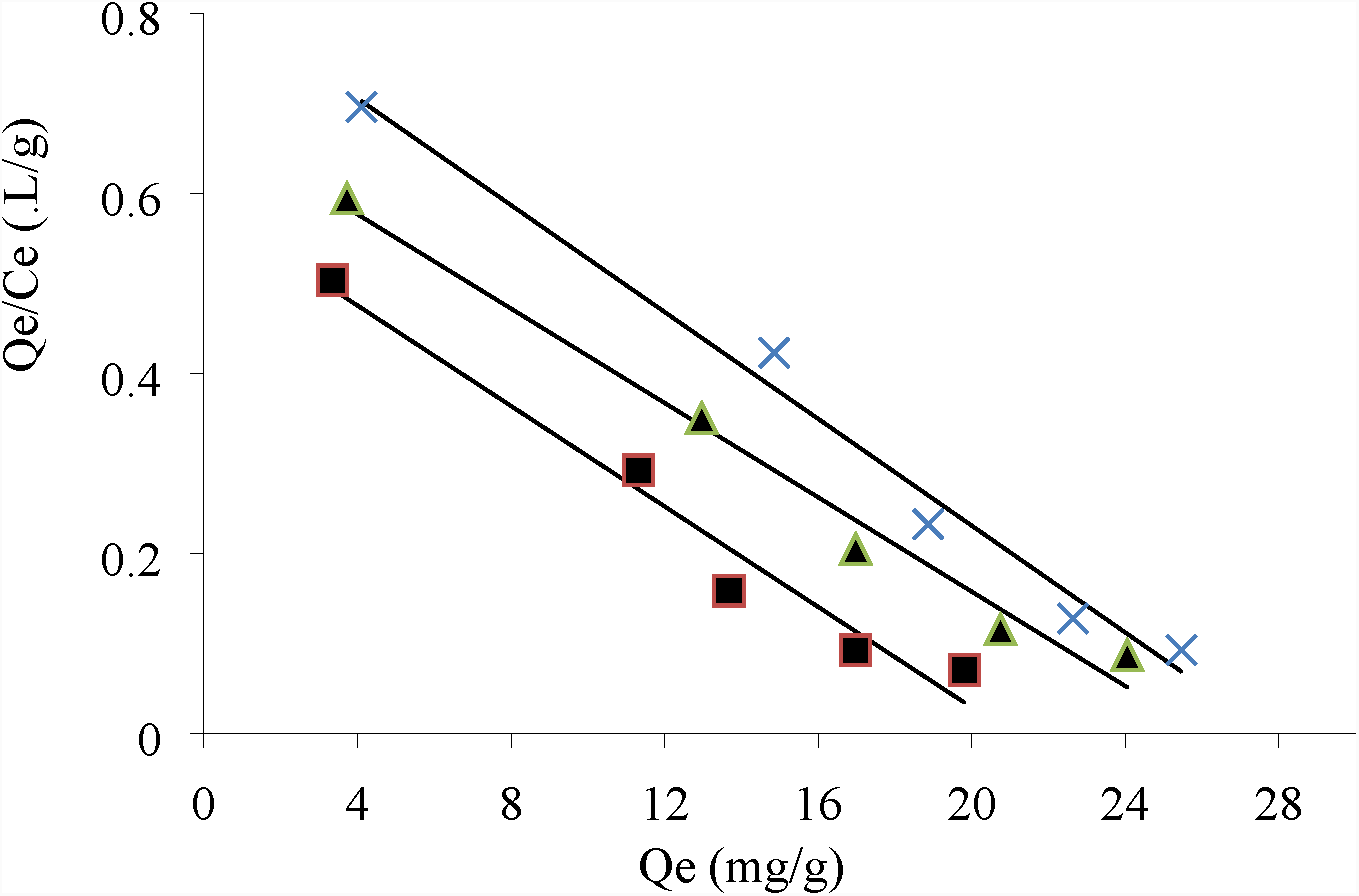
Scatchard plots for Cr(VI) ions onto *Eupatorium adenophorum*–alginate beads at initial Cr(VI) concentration from 10 to 300 mg/L, pH 2.0, contact time 60 min and biomass concentration 1.0 g/L at 20 (-▄--), 30 (-×-) and 40°C (-▴-) respectively.

The correlation coefficient values were found to be lower for Cr(VI) with comparison with Langmuir model, indicating that biosorption of Cr(VI) ions was not followed by Freundlich isotherm model. However, values of Freundlich constant, *n* in the range of 2 to10 suggested that Cr(VI) sorption is favorable on *Eupatorium adenophorum*-alginate beads [24].

The values of correlation coefficient of linear regression (*R*^*2*^) were found quite low for Gin isotherm model, suggesting that the model was unable to describe the Cr(VI) biosorption on *Eupatorium adenophorum*-alginate beads. However, the negative values of *β* confirmed the feasibility of Cr(VI) biosorption [25].

The relatively low values of *R*^*2*^ were obtained with equilibrium data, showing that Temkin model is not suitable for Cr(VI) biosorption on this biomass. Heat of biosorption, *b*_*T*_ was found to be positive at all temperatures, suggesting that Cr(VI) biosorption is exothermic in nature. In addition, lower values of *b*_*T*_ obtained in this study indicated the weak interactions between Cr(VI) ions and binding sites, revealing that physisorption as predominant mechanism [26].

Dubinin-Radushkevich (D-R) isotherm model was also unable to describe the experimental data as evidenced by the low correlation coefficient values. It was also found that the calculated *Q*_*m*_ values are much higher than experimental values at all temperatures. As it can be seen from Table 3, the numerical values of *E* suggested the physisorption is the predominant mechanism for Cr(VI) sorption on *Eupatorium adenophorum*-alginate beads [27].

Flory-Huggins model was also unable to correlate the experimental equilibrium data, due to lower correlation coefficients values. The results showed that Gibbs free energy values are negative, which indicated the spontaneous nature and feasibility of Cr(VI) sorption on *Eupatorium adenophorum*-alginate beads and the sorption is an exothermic reaction [4].

High correlation coefficient values indicated that Hill-der Boer model is good agreement with equilibrium data. The higher values of *k*_1_ and lower values of *k*_2_ indicated the strong interactions of Cr(VI) ions on *Eupatorium adenophorum* Sprengel surface binding sites [28]. It also showed that Cr(VI) ions are strongly interacted on surface binding sites at 30 °C compared to 20 and 40 °C respectively [4].

The calculated values of correlation coefficient by Hasely isotherm model are insignificant, confirming the unable of describing the experimental equilibrium data at tested temperatures. It also indicated that multilayer sorption is not involved in Cr(VI) biosorption on *Eupatorium adenophorum*-alginate beads [29].

### Thermodynamic analysis

The fundamental thermodynamic parameters of Cr(VI) removal using *Eupatorium adenophorum*-alginate beads at different temperature are presented in Table 5. The negative values of Δ*G°* obtained at all temperatures indicated the spontaneous nature of Cr(VI) biosorption. Horsfall et al., [38] suggested that Δ*G°* values up to −20 kJ/mol indicate the physical sorption, and those values greater than −40 kJ/mol suggest the chemical sorption. The calculated values of Δ*G°* indicated that Cr(VI) sorption on *Eupatorium adenophorum*–alginate beads proceeded with physical sorption. The negative value of Δ*H°* indicated the exothermic nature of biosorption process. Moreover, magnitude of Δ*H°* also gives information on biosorption nature, which may be either physical or chemical. According to Sag and Kustal, [39], Δ*H°* values ranging from 2.1 to 20.9 kJ/mol correspond to physisorption, whereas values ranging from 20.9 to 418.4 kJ/mol indicate the chemisorption. The value of Δ*H°* obtained in the range of 20 to 40 °C suggested the physisorption of Cr(VI) ions on calcium alginate entrapped *Eupatorium adenophorum* Sprengel biomass. The positive value of entropy change (Δ*S°*) indicated that the increased randomness at the solid/solution interface during the biosorption, suggesting that the affinity of metal ions on biomass surface.

**Table 5.** Thermodynamic parameters of Cr(VI) biosorption onto *Eupatorium adenophorum*-alginate beads

| $C_o$ (mg/L) | $T$ (°C) | Thermodynamic parameters | | | | |
| --- | --- | --- | --- | --- | --- | --- |
| | | $\Delta G^\circ$ (kJ mol<sup>-1</sup>) | $\Delta H^\circ$ (kJ mol<sup>-1</sup>) | $\Delta S^\circ$ (J mol<sup>-1</sup>K) | $E_a$ (kJ mol<sup>-1</sup>) | $\Delta H_r$ (J mol<sup>-1</sup>K) |
| 10 | 20-40 | -18.156 to-18.323 | 2.612 | 53.549 | 0.711 | -5.144 |

According to Ayoob et al., [31], *E*_a_ values in the range of 8 to 25 correspond to physisorption, less than 21 to aqueous diffusion, 20–40 to pore diffusion, whereas the values greater than 84 suggest the ion exchange as the sorption mechanism. The value of *E*_a_ obtained in the biosorption of Cr(VI) ions onto *Eupatorium adenophorum*-alginate beads may involve with weak interactions between the surface binding sites and Cr(VI) ions. Specifically, relatively low *E*_*a*_ value suggested that the biosorption has a low potential energy barrier.

It was found that Δ*H*_*r*_ values in the range of 20 to 40 °C, indicating that physisorption is involved in Cr(VI) biosorption with Ca-alginated *Eupatorium adenophorum* Sprengel biomass [32]. The positive value of Δ*Hr* suggested that Cr(VI) biosorption is exothermic in nature, which is in further compliance with the previous results derived from the estimation of thermodynamic parameters.

### Effect of interfering co-ions

The presence of co-ions including metal cations and anions in wastewater may cause interference and competition phenomena for biosorption sites. Table 6 shows the effect of co-existing ions on Cr(VI) sorption onto *Eupatorium adenophorum*-alginate beads. The results demonstrated that there is great selectivity towards Cr(VI) ions in the presence of Mg^+2^, Ca^+2^, Zn^+2^, Cd^+2^, Cu^+2^ and Ni^+2^, possibly due to their inability to chelate the surface binding sites. On the other hand, Cr(VI) sorption efficiency is slightly decreased with increase in concentration of SO^4-^, Cl^-^, and CO_3_^-^ ions respectively. It is due to the competition of negative ions in the same binding on the biomass surface. It was also found that sorption efficiency of Cr(VI) increases with increasing the concentration of Fe^+3^ ions. This may be attributed to the interactions between positively charged >FeOH_2_^+^ groups of biomass surface and HCrO_4_^-^ ions [40]. In another series of competitive experiments, a multi-ions system of 10 components was used in order to investigate their effect on primary target Cr(VI) ions. The removal efficiency was estimated at 28.72, 15.75 and 6.98% for Cr(VI) in the presence of 5, 10 and 25 mg/L of each ion used, suggesting their antagonistic action for the same binding sites on the biomass surface.

**Table 6.** Effect of interfering ions on Cr(VI) biosorption at initial concentration of 10 mg/L at optimum conditions

| Co-ions (mg/L) | Cr(VI) removal (%) |  |  |  |  |  |  |  |  |  |
| --- | --- | --- | --- | --- | --- | --- | --- | --- | --- | --- |
|  | SO<sub>4</sub><sup>-2</sup> | Cl<sup>-</sup> | CO<sub>3</sub><sup>-2</sup> | Mg<sup>+2</sup> | Ca<sup>+2</sup> | Fe<sup>+3</sup> | Zn<sup>+2</sup> | Cd<sup>+2</sup> | Cu<sup>+2</sup> | Ni<sup>+2</sup> |
| 5 | 41.10 | 43.53 | 43.08 | 38.18 | 38.22 | 43.08 | 41.08 | 44.55 | 41.09 | 36.01 |
| 10 | 40.03 | 38.18 | 40.58 | 38.03 | 39.81 | 46.77 | 41.75 | 38.99 | 38.50 | 38.55 |
| 25 | 41.61 | 38.97 | 41.12 | 40.89 | 41.38 | 51.04 | 39.65 | 42.05 | 40.28 | 39.11 |
| 50 | 37.78 | 36.71 | 36.97 | 42.66 | 38.74 | 59.33 | 40.47 | 42.13 | 42.00 | 42.67 |

### Desorption studies

The spent *Eupatorium adenophorum*-alginate beads, which contains Cr(VI) is unsafe for disposal due to stringent environmental regulations. The desorption of Cr(VI) ions from metal-loaded *Eupatorium adenophorum*-alginate beads was observed at 92.091%. This result indicated that Cr(VI) ions were adsorbed extracellularly by binding sites. The high desorption efficiency of Cr(VI) ions from Cr(VI)-loaded biomass within a short period of time suggested the weak interactions between Cr(VI) ions and biomass surface binding sites. Ranganathan, [33] reported the 65% of Cr(VI) desorption from the *Casurina equisetifolia* biomass using NaOH and HCl as desorbing agents.

## Conclusions

Calcium alginate entrapped *Eupatorium adenophorum* Sprengel stems powder biomass showed the high removal efficiency for removal of Cr(VI) species from aqueous solutions. It was found that sorption process was affected by the solution pH, biomass concentration, contact time, and initial Cr(VI) concentrations respectively. The results showed that the biosorption of Cr(VI) by *Eupatorium adenophorum Sprengel*-alginate beads followed the Langmuir isotherm model. The pseudo-second-order kinetics was found to explain the kinetics of biosorption more effectively based on the agreement of calculated and experimental uptake capacities along with high correlation coefficient values. Intraparticle diffusion model indicated that Cr(VI) biosorption may take place in two steps. It was also found that selectivity for Cr(VI) biosorption was observed in binary mixtures with various interfering ions. Thermodynamic parameters indicated the feasibility, spontaneous and exothermic nature of Cr(VI) biosorption. It can be concluded that *Eupatorium adenophorum* Sprengel-alginate beads can be proposed as an excellent biosorbent with potentially important applications in removal of heavy metals from contaminated sites.

